# Quantitative dialing of gene expression via precision targeting of KRAB repressor

**DOI:** 10.1101/2020.02.19.956730

**Authors:** Matthew S. Wilken, Christie Ciarlo, Jocelynn Pearl, Elaine Schanzer, Hanna Liao, Benjamin Van Biber, Konstantin Queitsch, Jordan Bloom, Alexander Federation, Reyes Acosta, Shinny Vong, Ericka Otterman, Douglass Dunn, Hao Wang, Pavel Zrazhevskiy, Vivek Nandakumar, Daniel Bates, Richard Sandstrom, Fyodor D. Urnov, Alister Funnell, Shon Green, John A. Stamatoyannopoulos

## Abstract

Human genes are regulated quantitatively, yet the ability to specify the expression level of a native gene accurately and specifically using a defined reagent has remained elusive. Here we show that precise targeting of KRAB repressive domain within regulatory DNA unlocks an endogenous quantitative ‘dial’ that can be engaged at nucleotide resolution to program expression levels across a wide physiologic range, with single-gene specificity and high reproducibly in primary cells.

## Main

In their native state, genes are regulated quantitatively to produce specific biological outcomes. Achieving such tunable gene expression is a key goal for mechanistic studies of gene function, therapeutic cell engineering, and synthetic biology. To date, no method has been described that provides single-gene-specific, incremental control of endogenous expression levels under uniform dosing conditions, particularly without requiring genome modification.

Most approaches to quantitative control of gene expression have relied on genomic integration of regulatory constructs^1-3^. A synthetic promoter can be placed under control of exogenous small molecules such as tetracycline to produce a quantitative range of gene expression^4-6^. MicroRNA elements can be recoded to tune gene expression^3^. While RNAi provides some degree of tunable repression without genome modification, it is plagued by variable efficacy and widespread off-target effects^7-10^.

In the context of dCas9, synthetic repressor activity can be modulated by small molecule control of RNA-guided delivery, but achieving defined expression levels is challenging^11^. Genomic targeting of dCas9-KRAB can be attenuated by engineering mismatched guide RNAs^12^, but this approach carries significant potential for untoward effects such as off-targeting.

Native transcription factors (TFs) convert information encoded in regulatory DNA regions such as promoters and enhancers into gene expression and cell state outcomes. TFs are modular proteins that combine a DNA recognition domain with one or more domains that confer specific functions via interplay with other chromatin-associated proteins^13, 14^. Coupling synthetic DNA binding domains with naturally-occurring KRAB repressor domains is a widely-applied approach for modulating gene expression, chiefly for the goal of gene silencing^15-20^. KRAB domains recruit the KAP1 co-repressor and, in turn, endogenous enzymatic complexes that methylate histones and DNA and trigger focal heterochromatin formation^15-20^. Despite decades of work, however, it remains unclear what factors contribute to KRAB activity in the context of a given proximal regulatory region.

Regardless of the DNA targeting modality employed, observed potencies of synthetic KRAB repressors have been highly variable, and reliably achieving complete repression comparable to gene knockout has been particularly elusive^7, 21-23^. KRAB also has the potential to trigger mitotically heritable gene repression^24-27^, yet its application for this purpose has likewise been confounded by variable effects depending on experimental context and gene targeted^17, 24, 27-30^.

Here we report a generalizable approach for achieving quantitative, highly specific, and heritable gene expression states in primary cells. We demonstrate that KRAB repressor activity is dominantly dependent on the precise genomic position to which it is targeted, providing both a framework for achieving potent, durable repression of endogenous genes and an explanation for previously reported discrepancies in KRAB activity. We show that synthetic repressors targeted to nucleotides that gate near-complete abrogation of gene expression do so with single-gene specificity and can be readily multiplexed, opening new avenues for precision programming of genes and cells for both basic and therapeutic applications.

## Results

### Nucleotide-precise delivery of KRAB repressor domains to endogenous promoters

To achieve nucleotide-precise targeting of KRAB domains to specific promoter positions, we utilized *Xanthomonas* TAL effector repeats which enable modular synthesis of DNA binding domains (DBDs) capable of targeting ∼95% of the human genome sequence^31, 32^. Synthetic TAL DBDs (T-DBDs) can be appended at either their C-or N-termini with effector domains conferring function in mammalian cells, for example the KRAB repressor domain^24, 29, 33-36^.

As a test case, we focused on a well-characterized immune checkpoint gene *TIM3 (HAVCR2)*, which encodes a cell surface molecule that can be robustly quantified by flow cytometry. We designed a series of densely spaced synthetic T-DBD-KRAB repressors targeting the TIM3 promoter (Fig. 1A, top). To quantify potency, we electroporated each repressor mRNA into primary CD3+ T cells and measured surface expression of TIM3 after 48 hours.

**Fig. 1.**
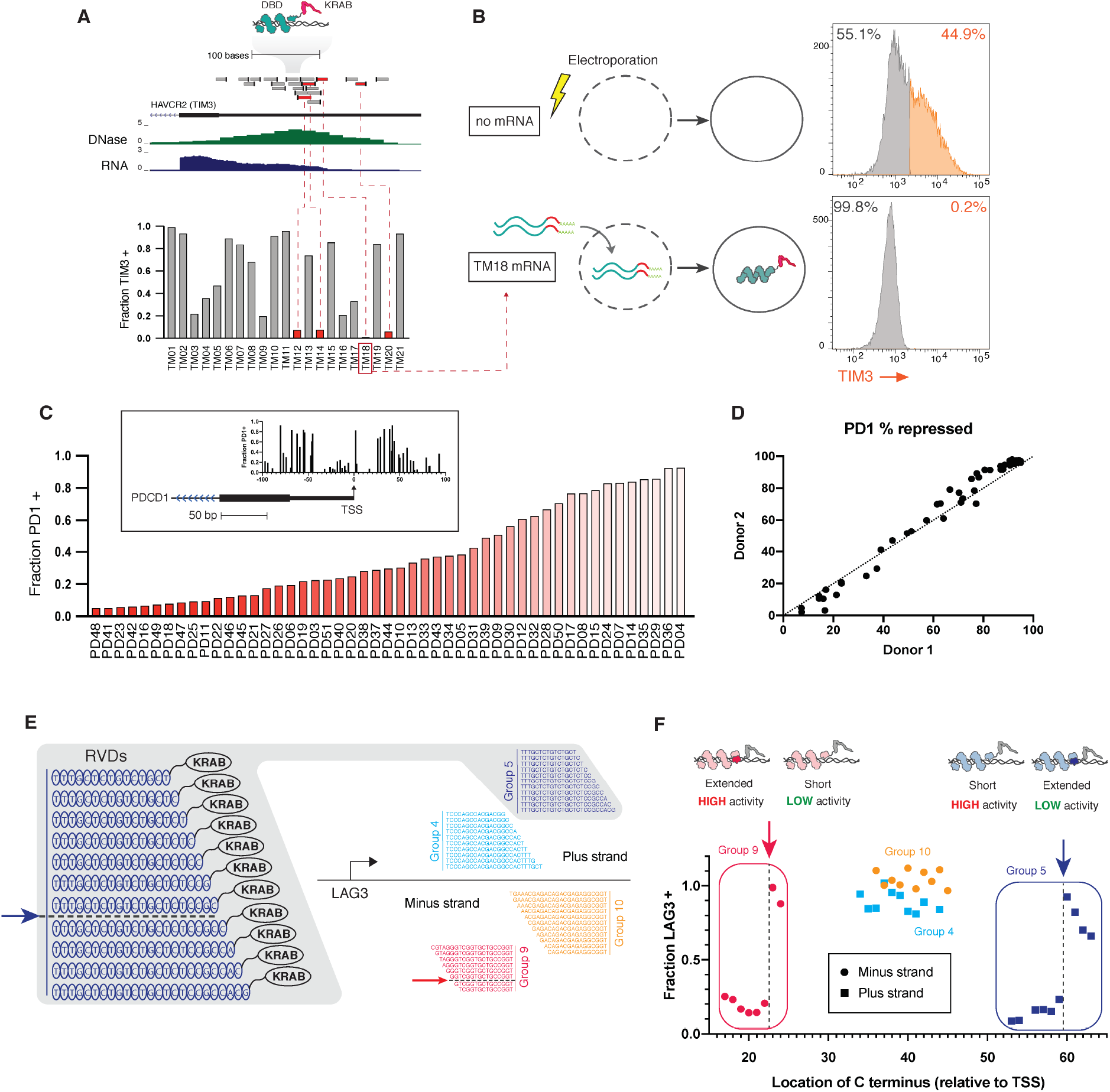
Quantitative repression achieved by nucleotide-precise targeting of KRAB to promoter DNA. (a) Selection of repressors at the *TIM3 (HAVCR2)* promoter. Top: DNA binding domains (grey boxes) are shown to scale; tick marks indicate position of C-terminal KRAB domain. Red indicates ‘keyhole’ sites: DBD-KRABs with >90% repression. Center: DNase-seq and RNA-seq normalized tag density. Bottom: Fraction of TIM3+ cells (normalized to mock transfection) as quantified by flow cytometry 48 hours after electroporation of repressor mRNA into activated CD3+ human T-cells. (b) TIM3 surface protein expression quantified by flow cytometry at 48 hours post-transfection (plots representative of three independent experiments). (c) SynTFs targeting different locations in the *PD-1 (PDCD1)* promoter produce a finely graded range of repression levels. Fraction PD-1+ cells (normalized to mock transfection) was quantified by flow cytometry 48 hours after electroporation of repressor mRNA into activated CD3+ human T-cells. Inset shows repressor activity as a function of position of C-terminal KRAB domain. (d) Percent repression of PD-1 in two independent experiments using CD3+ T cells from two different donors. Each point represents an individual synTF from (c). Dotted line x=y shown for reference. (e) T-DBDs targeting seed sequences in the *LAG3* promoter were sequentially extended by one repeat unit to produce groups of T-DBD-KRABs with different positioning of the C-terminal KRAB (see example group). (f) Fraction LAG3+ cells at 2 days post-transfection as measured by flow cytometry, normalized to no RNA controls. X-axis indicates the location of the KRAB domain relative to the *LAG3* TSS. Groups 5 and 9 are highlighted to demonstrate loss/gain of repression activity when the KRAB domain was moved by one nucleotide.

Varying the genomic positioning of T-DBDs produced a quantitative landscape of gene expression (Fig. 1A, bottom). *A priori*, we expected that repressors targeted with close proximity would be nearly equivalent in function. Instead, we found synthetic repressor activity was highly variable even between closely spaced repressors. Within this landscape, we observed a small subset of positions that yielded dramatic drop-offs in gene expression, resulting in near-complete repression (Fig. 1B). We termed such positions ‘keyhole’ sites for repression. Repressors targeting keyhole sites produced near-complete gene silencing, which was accompanied by loss of H3K4me3 and gain of H3K9me3 as expected for KRAB-induced silencing (Fig. S1).

To examine extensibility and quantitative reproducibility of expression levels programmed by positional targeting of KRAB, we tiled T-DBD-KRABs near the transcription start site of *PD-1* (*PDCD1*) and quantified PD-1 expression in CD3+ T cells 48 hours after repressor mRNA electroporation. We observed a similar quantitative spectrum of repression, including highly active keyhole sites, spanning the entire range of physiologic PD-1 expression (Fig. 1C). Next, we repeated the experiment using the same set of T-DBD-KRABs delivered to an independently collected and temporally separated T cell sample from a different donor. Position-specific repression levels were highly reproducible between donors and experiments, demonstrating the robust incremental expression control achievable by targeting specific KRAB to specific genomic positions (Fig. 1D).

### A single nucleotide positional trigger for KRAB-induced repression

The precipitous differences in repression we observed as a function of genomic position suggested that the triggering of repression by KRAB might be under very fine positional control. To test this, we devised a strategy for migrating a KRAB domain at 1 bp intervals by incrementally extending DNA binding domains anchored from a common 5’ position (Fig. 1E). We synthesized a total of 40 T-DBD-KRAB repressors extending from 4 anchor points, providing per-base coverage of 40 nucleotide positions across both strands of a region within the *LAG3* promoter encompassing two positions where repressor activity was identified in an initial screen (Fig. 1E). Quantification of LAG3 levels from each positional variant individually in primary CD3+ T cells revealed discrete positions where migrating the KRAB domain even 1 bp 3’ or 5’ was sufficient to trigger strong repression from otherwise identical T-DBD-KRAB molecules (Fig. 1F). Repressor activity did not correlate with genomic features such as DNA accessibility or distance from the transcription start site of a gene (Fig. S2). Furthermore, there was no apparent dependence of repressor activity on DBD length, as would be expected if DBD affinity or residence time were the main determinant of activity^37-39^ (Fig. 1F, Fig. S3). Our results indicate that the epigenetic silencing cascade initiated by KRAB is precisely triggered at single nucleotide resolution reflecting its linear (and hence rotational) positioning within promoter chromatin.

### Potent repressors are single gene-specific

Potency is often accompanied by off-target effects or toxicity. We therefore sought to quantify the specificity of highly potent repressors for their genic targets by RNA-seq, a sensitive measure of both on- and off-target effects genome-wide. We delivered potent keyhole repressors of the immune checkpoint genes *TIM3, LAG3*, and *PD-1* to primary CD3+ T cells both individually and simultaneously as a pool and performed total RNA-seq at 48h when peak repression is achieved (Fig. 2A-C left, genome browser views). Individual repressors ablated RNA expression of their target genes with near complete specificity (Fig. 2A-C right, volcano plots). Of note, the *LAG3* repressor resulted in down-regulation of the closely positioned gene *PTMS* located ∼1kb upstream (Fig. 2B), consistent with a +/-∼2kb H3K9me3 ‘halo’ produced by KRAB-triggered silencing (Fig. S1). While *LAG3* was completely repressed, *PTMS* was only partially repressed (35% of control) (Fig. 2B); both are on-target effects of the same target site.

**Fig. 2.**
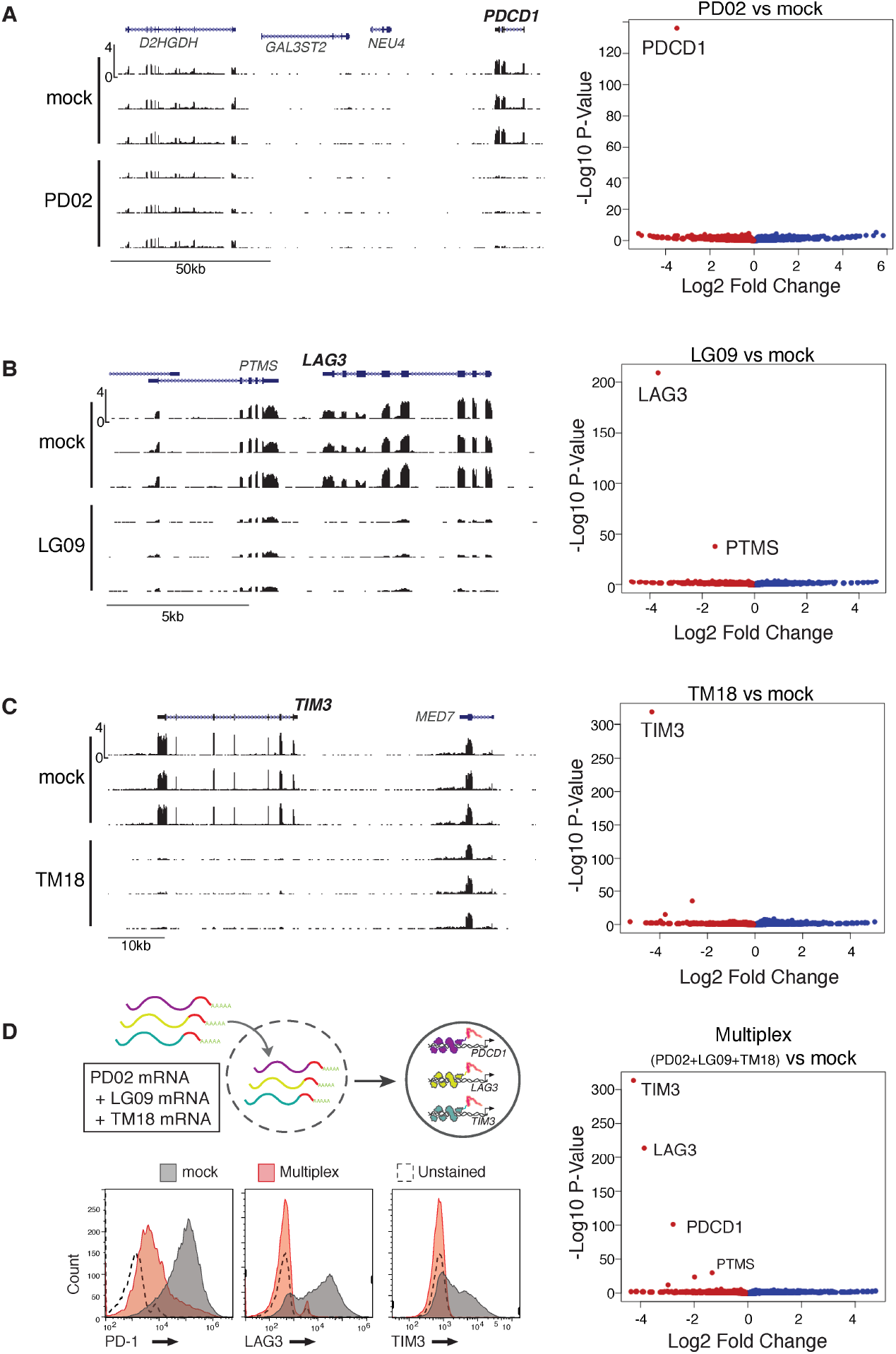
High specificity of keyhole repressors singly and in multiplex. (a) <u>Left</u>: RNA-seq tag density (scale bar in upper left) in the *PDCD1 (PD-1)* locus following blank electroporation (top tracks, three independent replicates), and delivery of synthetic repressor PD02 targeting *PDCD1* (bottom tracks, three independent replicates). <u>Right</u>: Volcano plot showing differential gene expression (RNA-seq q-value, vertical axis) following PD02 repressor delivery. (b) Results for *LAG3* synthetic repressor, as for (a). (c) Results for *TIM3* repressor, as for (a, b). Note that in addition to complete repression of *LAG3*, the *PTMS* gene located 1.2kb upstream is partly repressed. (d) Left, co-delivery of PD02, LG09, and TM18 results in repression of target genes similar to individually delivered repressors. Right, volcano plot of differential gene expression (RNA-seq) q-values for co-delivered repressors consistent with linearly additive (and independent) contribution of each repressor.

Multiplexing provides an even more stringent test of specificity and effector independence. Simultaneous delivery of all three repressors produced purely additive effects, with no loss of potency or specificity (Fig. 2D). We also observed both additivity and dose-dependence at the level of a single gene targeted by multiple synthetic repressors directed to different sites within the same promoter (Fig. S4). Taken together, these results indicate that even highly potent synthetic repressors exhibit remarkable specificity whether delivered individually or in multiplex.

### Transient KRAB-induced repression is reliably mitotically heritable

We next studied the duration of transcriptional repression as a function of synthetic repressor persistence. Repressor mRNA and protein are rapidly degraded following electroporation, with protein returning to background levels by 48h post mRNA electroporation as measured by direct immunofluorescence (Fig. 3A-B). Following CD3/CD28 stimulation, primary T cells begin cycling with a doubling time of approximately 36 hours (Fig. S5). As such, effects on gene expression persisting beyond 72 hours reflect mitotically heritable states. In mock transfected cells, TIM3 expression peaks at 8 days post stimulation before beginning a gradual decline to steady state levels of ∼40% TIM3+ cells (Fig. 3C, open circles). By contrast, cells receiving the TM18 repressor show near complete repression of *TIM3* up to day 5 post electroporation (day 7 post stimulation) and persistent repression in a declining subpopulation of cells for another ∼20 days, the practical limit of T cell culture (Fig. 3C, solid black circles, red trace). Even more pronounced longitudinal repression was induced by a synthetic repressor targeting *PD-1* and persisted for approximately two weeks in culture (Fig. 3D). These results show that potent repression by positionally-targeted KRAB is mitotically heritable, with variable multi-day kinetics observed for different genes.

**Fig. 3.**
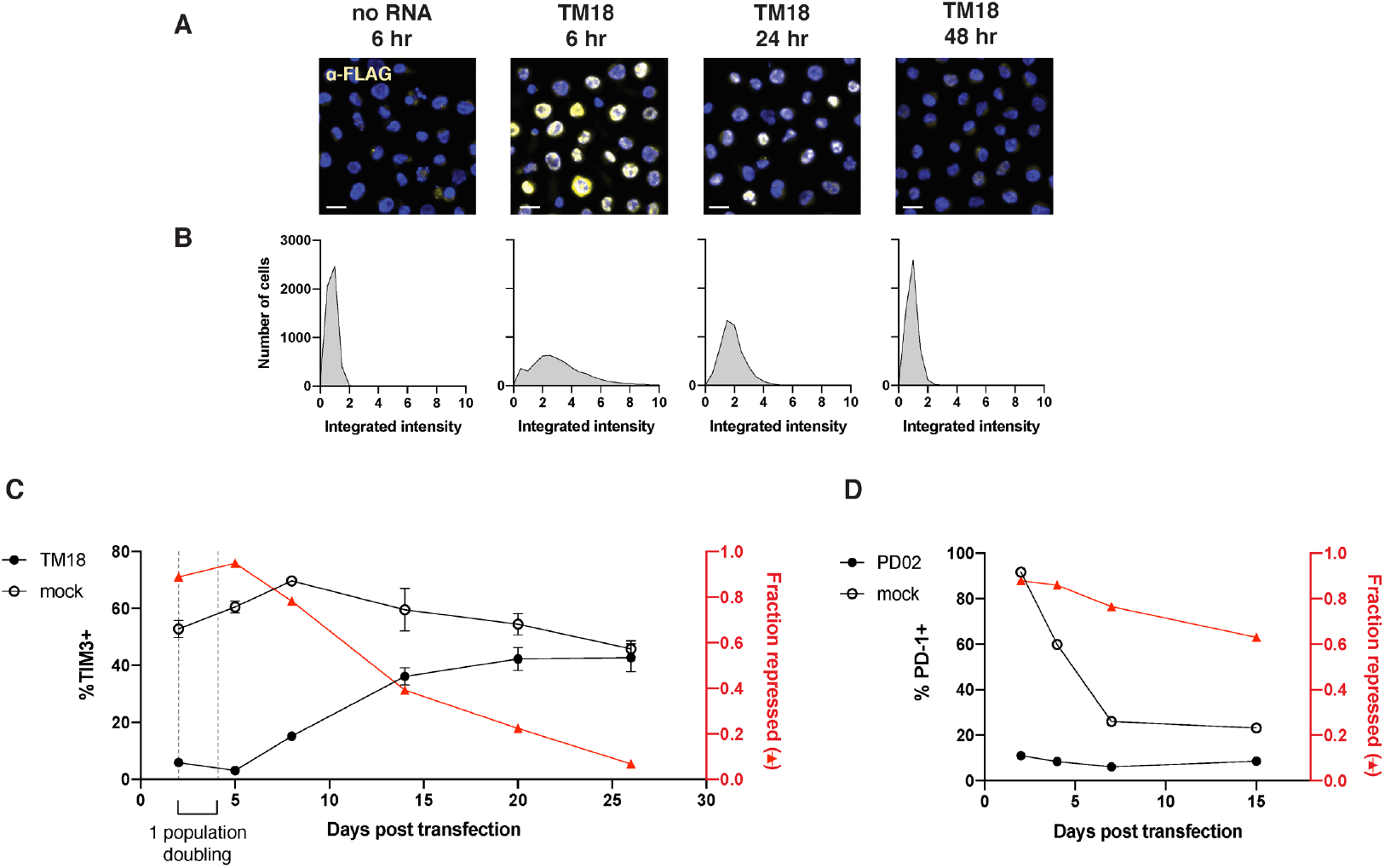
Mitotically heritable repression produced by targeting keyhole sites. (a) Expression of repressor protein over time. Cells were electroporated with either no RNA or the *TIM3* repressor TM18 at time 0, and TM18 protein levels were determined by anti-FLAG immunofluorescence (yellow) for up to 48 hours. Nuclear DAPI staining is shown in blue. Scale bar indicates 10 um. (b) Histograms show integrated anti-FLAG fluorescence intensity per nucleus over a population of cells with a bin size of 0.5. (c) Kinetics of *TIM3* repression by synthetic repressor TM18. Primary T cells were electroporated with either no RNA or TM18 at day 0, and TIM3 expression was determined by cell surface antibody staining and flow cytometry for 26 days. Cells with greater fluorescence intensity than unstained control were considered TIM3+. Percent TIM3+ cells is indicated by dark colored lines and left y-axis. Fraction of cells with *TIM3* repressed (%TIM3 negative cells in TM18-treated cells relative to no RNA control) is indicated by the right y-axis and red line. Bars indicate standard deviation of two electroporations in the same experiment. Repression was maintained through a time period equivalent to one population doubling. Population doubling time was calculated assuming a constant proliferation rate and cell counts at days 2 and 5. (d) Kinetics of *PD-1* repression by synthetic repressor PD02 as in (a). PD-1 expression was determined over the course of 15 days post-electroporation.

## Discussion

Human genes are regulated quantitatively, and the ability to specify their expression level using defined reagents would have broad applications in biology and therapeutics^40^. Our results show that the precise genomic position within the proximal regulatory region of an endogenous human gene quantitatively specifies the level of repression produced by a KRAB repressor domain targeted to that position, with some positions conferring near-complete repression. These effects are independent of DBD length (and hence affinity and residence time^37-39^), affirming the dominant contribution of genomic position. Position-specific expression levels are quantitative over a wide range and are highly reproducible, providing an endogenous genomic ‘dial’ that can be turned to deliver a desired expression level with true single-target specificity. Notably, this level of functional specificity has not been reported with other editing modalities^24, 29, 41-43^.

Beyond a general methodology for programming gene expression, our results also provide a unifying explanation for the widely variable and sometimes contradictory results obtained to date using synthetic KRAB-containing repressors. Both the level and durability of repression induced by KRAB has been reported to vary widely from gene to gene, even when the same types of constructs are employed^7, 22^-^24, 29, 30^, suggesting that the KRAB domain might need to be combined with additional functional domains in order to obtain potent or heritable repression^23^. However, our results show that this is not the case.

Like the DBDs of endogenous transcription factors, TALE DBDs engage the genome in its native double-stranded form, in contrast to RNA-guided protein-DNA recognition by Cas9, which involves extensive unwinding and disruption of the DNA template^44, 45^. While some screening studies have implicitly incorporated low-resolution positional targeting of dCas9-KRAB^46-48^, this has invariably been in the context of pooled experiments with enrichment-based readouts that lack quantitative information about gene expression levels per guide tested. Moreover, any observed positional variability in dCas9-KRAB-induced repression must be corrected for nucleosome occupancy which has a dominant effect on dCas9 engagement^46, 47^.

Recent studies have reported both naturally-occurring and synthetic KRAB variants with increased intrinsic potency relative to conventional KRAB^21, 49^. We note that the reported relative repressive contribution of novel KRAB variants is typically considerably smaller than the wide dynamic range conferred by nucleotide-positional targeting. As such, an enhanced or attenuated KRAB domain would be expected to be dominated by position-dependence, though would offer a strategy for further enhancing or attenuating position-specified effects. Our results thus suggest that any future studies of the impact of variant KRAB domains or the combination of KRAB with additional functional domains on gene expression levels and mitotic heritability should thoroughly account for position dependence.

The observed strict dependence of repression on genomic position suggests a structural mechanism under which a specific positional/rotational presentation of the KRAB domain is necessary to successfully recruit KAP1 and trigger its sequelae. However, despite dramatic progress in structural biology, a detailed understanding of the biophysical architecture of even a single human regulatory region remains elusive^50^. Irrespective of the underlying mechanism, quantitative positional specification of repressive function should have broad applications in the engineering of endogenous and synthetic gene expression programs.

## Acknowledgements

We thank J. Halow and K. Lee for assistance with cell culture; M. Diegel and F. Neri for assistance with sequencing; J. Lazar for input on statistical analysis.

## Author contributions

M.S.W., C.C., J.P., A.F., F.D.U., S.G. and J.A.S. designed the research. M.S.W., C.C., J.P., E.S., J.B., H.L., B.V.B., K.Q, A.F., R.A., S.V., E.O., and A.F. performed cell engineering experiments. D.D., H.W., and D.B. performed RNA-seq and CUT&RUN experiments. P.Z. and V.N. performed imaging experiments. M.S.W., C.C., J.P., J.B., R.S., P.V., and V.N. analyzed data. M.S.W., C.C., S.G., and J.A.S. wrote the manuscript with input from other co-authors.

## Data and materials availability

All RNA-seq and imaging data, software code used for analysis, protein sequences, protocols, and materials used in the experiments and data analysis will be made freely available.

## References

1. Deans, T.L., Cantor, C.R. & Collins, J.J. A tunable genetic switch based on RNAi and repressor proteins for regulating gene expression in mammalian cells. Cell 130, 363–372 (2007).

2. Li, X.T. et al. tCRISPRi: tunable and reversible, one-step control of gene expression. Sci Rep 6, 39076 (2016).

3. Michaels, Y.S. et al. Precise tuning of gene expression levels in mammalian cells. Nat Commun 10, 818 (2019).

4. Gossen, M. & Bujard, H. Tight control of gene expression in mammalian cells by tetracycline-responsive promoters. Proc Natl Acad Sci U S A 89, 5547–5551 (1992).

5. Gossen, M. et al. Transcriptional activation by tetracyclines in mammalian cells. Science 268, 1766–1769 (1995).

6. Yin, D.X., Zhu, L. & Schimke, R.T. Tetracycline-controlled gene expression system achieves high-level and quantitative control of gene expression. Anal Biochem 235, 195–201 (1996).

7. Evers, B. et al. CRISPR knockout screening outperforms shRNA and CRISPRi in identifying essential genes. Nat Biotechnol 34, 631–633 (2016).

8. Jackson, A.L. et al. Expression profiling reveals off-target gene regulation by RNAi. Nat Biotechnol 21, 635–637 (2003).

9. Scacheri, P.C. et al. Short interfering RNAs can induce unexpected and divergent changes in the levels of untargeted proteins in mammalian cells. Proc Natl Acad Sci U S A 101, 1892–1897 (2004).

10. Semizarov, D. et al. Specificity of short interfering RNA determined through gene expression signatures. Proc Natl Acad Sci U S A 100, 6347–6352 (2003).

11. Gangopadhyay, S.A. et al. Precision Control of CRISPR-Cas9 Using Small Molecules and Light. Biochemistry 58, 234–244 (2019).

12. Jost, M. et al. Titrating gene expression using libraries of systematically attenuated CRISPR guide RNAs. Nat Biotechnol 38, 355–364 (2020).

13. Brent, R. & Ptashne, M. A eukaryotic transcriptional activator bearing the DNA specificity of a prokaryotic repressor. Cell 43, 729–736 (1985).

14. Lambert, S.A. et al. The Human Transcription Factors. Cell 172, 650–665 (2018).

15. Bellefroid, E.J., Poncelet, D.A., Lecocq, P.J., Revelant, O. & Martial, J.A. The evolutionarily conserved Kruppel-associated box domain defines a subfamily of eukaryotic multifingered proteins. Proc Natl Acad Sci U S A 88, 3608–3612 (1991).

16. Friedman, J.R. et al. KAP-1, a novel corepressor for the highly conserved KRAB repression domain. Genes Dev 10, 2067–2078 (1996).

17. Groner, A.C. et al. The Kruppel-associated box repressor domain can induce reversible heterochromatization of a mouse locus in vivo. J Biol Chem 287, 25361–25369 (2012).

18. Margolin, J.F. et al. Kruppel-associated boxes are potent transcriptional repression domains. Proc Natl Acad Sci U S A 91, 4509–4513 (1994).

19. Nawrath, M., Pavlovic, J. & Moelling, K. Inhibition of human hematopoietic tumor formation by targeting a repressor Myb-KRAB to DNA. Cancer Gene Ther 7, 963–972 (2000).

20. Vissing, H., Meyer, W.K., Aagaard, L., Tommerup, N. & Thiesen, H.J. Repression of transcriptional activity by heterologous KRAB domains present in zinc finger proteins. FEBS Lett 369, 153–157 (1995).

21. Alerasool, N., Segal, D., Lee, H. & Taipale, M. An efficient KRAB domain for CRISPRi applications in human cells. Nat Methods 17, 1093–1096 (2020).

22. Gilbert, L.A. et al. CRISPR-mediated modular RNA-guided regulation of transcription in eukaryotes. Cell 154, 442–451 (2013).

23. Yeo, N.C. et al. An enhanced CRISPR repressor for targeted mammalian gene regulation. Nat Methods 15, 611–616 (2018).

24. Amabile, A. et al. Inheritable Silencing of Endogenous Genes by Hit-and-Run Targeted Epigenetic Editing. Cell 167, 219–232 e214 (2016).

25. Ayyanathan, K. et al. Regulated recruitment of HP1 to a euchromatic gene induces mitotically heritable, epigenetic gene silencing: a mammalian cell culture model of gene variegation. Genes Dev 17, 1855–1869 (2003).

26. Bintu, L. et al. Dynamics of epigenetic regulation at the single-cell level. Science 351, 720–724 (2016).

27. Gjaltema, R.A.F. et al. KRAB-Induced Heterochromatin Effectively Silences PLOD2 Gene Expression in Somatic Cells and is Resilient to TGFbeta1 Activation. Int J Mol Sci 21 (2020).

28. Mandegar, M.A. et al. CRISPR Interference Efficiently Induces Specific and Reversible Gene Silencing in Human iPSCs. Cell Stem Cell 18, 541–553 (2016).

29. Mlambo, T. et al. Designer epigenome modifiers enable robust and sustained gene silencing in clinically relevant human cells. Nucleic Acids Res 46, 4456–4468 (2018).

30. O’Geen, H. et al. Ezh2-dCas9 and KRAB-dCas9 enable engineering of epigenetic memory in a context-dependent manner. Epigenetics Chromatin 12, 26 (2019).

31. Boch, J. et al. Breaking the code of DNA binding specificity of TAL-type III effectors. Science 326, 1509–1512 (2009).

32. Moscou, M.J. & Bogdanove, A.J. A simple cipher governs DNA recognition by TAL effectors. Science 326, 1501 (2009).

33. Cong, L., Zhou, R., Kuo, Y.C., Cunniff, M. & Zhang, F. Comprehensive interrogation of natural TALE DNA-binding modules and transcriptional repressor domains. Nat Commun 3, 968 (2012).

34. Maeder, M.L. et al. Targeted DNA demethylation and activation of endogenous genes using programmable TALE-TET1 fusion proteins. Nat Biotechnol 31, 1137–1142 (2013).

35. Mendenhall, E.M. et al. Locus-specific editing of histone modifications at endogenous enhancers. Nature Biotechnology 31, 1133–1136 (2013).

36. Zhang, Z., Wu, E., Qian, Z. & Wu, W.S. A multicolor panel of TALE-KRAB based transcriptional repressor vectors enabling knockdown of multiple gene targets. Sci Rep 4, 7338 (2014).

37. Clauss, K. et al. DNA residence time is a regulatory factor of transcription repression. Nucleic Acids Res 45, 11121–11130 (2017).

38. Geiger-Schuller, K., Mitra, J., Ha, T. & Barrick, D. Functional instability allows access to DNA in longer transcription Activator-Like effector (TALE) arrays. Elife 8 (2019).

39. Rinaldi, F.C., Doyle, L.A., Stoddard, B.L. & Bogdanove, A.J. The effect of increasing numbers of repeats on TAL effector DNA binding specificity. Nucleic Acids Res 45, 6960–6970 (2017).

40. Keren, L. et al. Massively Parallel Interrogation of the Effects of Gene Expression Levels on Fitness. Cell 166, 1282–1294 e1218 (2016).

41. Grimmer, M.R. et al. Analysis of an artificial zinc finger epigenetic modulator: widespread binding but limited regulation. Nucleic Acids Res 42, 10856–10868 (2014).

42. Stojic, L. et al. Specificity of RNAi, LNA and CRISPRi as loss-of-function methods in transcriptional analysis. Nucleic Acids Res 46, 5950–5966 (2018).

43. Thakore, P.I. et al. Highly specific epigenome editing by CRISPR-Cas9 repressors for silencing of distal regulatory elements. Nat Methods 12, 1143–1149 (2015).

44. Deng, D. et al. Structural basis for sequence-specific recognition of DNA by TAL effectors. Science 335, 720–723 (2012).

45. Nishimasu, H. et al. Crystal structure of Cas9 in complex with guide RNA and target DNA. Cell 156, 935–949 (2014).

46. Horlbeck, M.A. et al. Compact and highly active next-generation libraries for CRISPR-mediated gene repression and activation. Elife 5 (2016).

47. Horlbeck, M.A. et al. Nucleosomes impede Cas9 access to DNA in vivo and in vitro. Elife 5 (2016).

48. Gilbert, L.A. et al. Genome-Scale CRISPR-Mediated Control of Gene Repression and Activation. Cell 159, 647–661 (2014).

49. Tycko, J. et al. High-Throughput Discovery and Characterization of Human Transcriptional Effectors. Cell 183, 2020–2035 e2016 (2020).

50. Panne, D., Maniatis, T. & Harrison, S.C. An atomic model of the interferon-beta enhanceosome. Cell 129, 1111–1123 (2007).

