## Supplemental Materials for "Quantitative dialing of gene expression via precision targeting of KRAB repressor"

**This PDF file includes:**

Materials and Methods
Figs. S1 to S5
Supplementary Figure Legends
Table S1

### Materials and Methods

#### T-Cell Culture

T-cells were sourced from Vitalant (https://www.vitalant.org/Home.aspx) as either TRIMA LRS (Leukocyte Reduction System) chambers or TRIMA Residuals from TRIMA Apheresis Collection Kits (TerumoBCT; https://www.terumobct.com/trima). CD3+ cells were isolated from donor blood by negative selection using the EasySep™ Human T Cell Isolation Kit (STEMCELL Technologies, 17951) and cryopreserved. Upon thawing, cells were activated with Dynabeads™ Human T-Activator CD3/CD28 for T Cell Expansion and Activation (Gibco, 1132D) at a ratio of 1:1 beads:cells and cultured in T cell media consisting of 10% Heat Inactivated Fetal Bovine Serum (HyClone, SH30071.03) and 1% PenStrep (Corning, 30-002-CI) in RPMI 1640 media (HyClone, SH30027.02) passed through a 0.22 um filter and supplemented with recombinant human IL-2 (Peprotech, 200-02) at a final concentration of 200U/mL. Cells were cultured in 37°C 5% $CO_2$ humidified HeraCell incubators (Thermo Scientific). Cell counts were performed by 0.4% trypan blue dye exclusion (Invitrogen, T10282). T cell media supplemented with rhIL-2 was refreshed every 2-3 days and cell densities were maintained between 500,000-1.5 million cells/mL.

#### Synthetic Repressor Design and Assembly

TAL monomers were cloned and assembled into full length TALs with modifications to established methods[51, 52] into a pVAX-based plasmid and included an N-terminal 3x-FLAG tag and SV40 nuclear localization signal (sequences in table S1). Functional domains were selected by literature search for evidence of transcriptional repressive function and annotated DNA-binding domains removed *in silico* before synthesis and incorporation into TAL or heterodimer constructs. Functional domains were added by Infusion cloning (Takara Bio; catalog # 638909) onto the C-terminal end of the TAL. Functional domain constructs contained a 15 amino acid linker domain (GGGGGMDAKSLTAWS) and a KRAB domain derived from human KOX1 (RTLVTFKDVFVDFTREEWKLLDTAQQIVYRNVMLENYKNLVSLGYQLTKPDVILRLE KGEEP).

#### *In vitro* Transcription of Synthetic Repressors

mRNA was generated by *in vitro* transcription (IVT) using the T7 mScript™ Standard mRNA Production System (CELLSCRIPT, C-MSC100625), including 5′-capping and poly-A addition reactions. RNA quality control was performed on a Fragment Analyzer Infinity (Agilent) using the standard RNA kit (DNF-471-1000). RNA was quantified based on absorbance at 260 nm. TALE plasmids were digested with Eag1 and column purified prior to IVT. Synthetic DNA was PCR amplified prior to IVT using the KAPA HiFi HotStart ReadyMix (Roche, KK2601) and column purified prior to IVT (Zymo, D4033).

#### Repressor Delivery to T Cells via Electroporation

Two days after thawing and activation (unless otherwise indicated), CD3+ T cells were spun down at 400xg for 5 minutes, then washed with PBS (Corning, 21-040-CM). Cells were resuspended in the appropriate volume of BTXpress High Performance Electroporation Solution (BTX, 45-0805) to give 250,000 cells per 100 uL electroporation. Cells were multi-channel pipetted into a PCR plate loaded with 1ug *in vitro* transcribed mRNA (for all repressors and heterodimer components) and mixed gently, then transferred to the MOS 96-Multi-Well Electroporation Plate_2mm (BTX, 45-0450).

Electroporation was performed using the ECM 830 Square Wave Electroporation System with 96-well HT-200 plate handler at 250 V for 5 ms. Post-electroporation, cells were transferred to a 96-well deep well plate (Axygen, P-DW-11-C-S) with 800 uL of warm T cell media + rhIL-2 and electroporation plate wells were rinsed with media.

Flow Cytometry

Cells were multi-channel pipetted into a 96-well V-bottom plate (Corning, 3894) and spun down (all spins at 500xg for 4 minutes). Cells were washed and spun once with 1X PBS (Corning, 21-040-CM) before staining with fluorophore-conjugated antibodies diluted 1:50 in 1X PBS for 30 minutes in the dark at room temperature. Following staining, FACS buffer (2% Heat-inactivated Fetal Bovine Serum (HyClone, SH30071.03), 1 mM EDTA (OmniPur, 4050) in 1X PBS, passed through a 0.22 um filter) was added to the samples and spun down. Cells were washed and spun down once more in FACS buffer before a final resuspension in FACS buffer for flow cytometry analysis. Antibodies used were: Brilliant Violet 421™ anti-human CD366 (Tim-3) Antibody (BioLegend, 345008); PE/Dazzle™ 594 anti-human CD223 (LAG-3) Antibody (BioLegend, 369332); APC anti-human CD279 (PD-1) Antibody (BioLegend, 329908); PE anti-EGFR Antibody (BioLegend 352904); AlexaFluor700 anti-human CD3 Antibody (BioLegend 300424). Flow analysis was performed on CytoFlex S (Beckman Coulter, B75442). Gating using unstained pooled cells or no mRNA controls.

RNA-seq data collection

RNA was collected from 5E5 to 1E6 cells 48-hours post electroporation, washed twice with PBS, pelleted at 500g for 5-minutes, resuspended in 350uL Buffer RLT (Qiagen), and frozen at -80C. RNA was isolated with Qiagen's RNeasy Micro kit (74004) with on column DNase treatment (RNase-Free Dnase Set #79254). RNA QC\quantification was performed on a Fragment Analyzer (Agilent) using kit # DNF-471-1000 (Standard RNA). RNA libraries were generated using Illumina's Truseq Stranded Total RNA-HT (with Ribo-Zero Gold) kit, #20020599, library QC\quantification was performed on our Fragment Analyzer (Agilent) using kit # DNF-474-1000 (High Sensitivity NGS). Sequencing was performed at 2x76bp on Illumina's Hiseq 4000.

RNA-seq data analysis

Raw sequencing reads were trimmed to remove adapter sequences and aligned to the human genome (hg38/GRCh38, no alts). RNA read mapping informatics was performed using RNA-STAR (2.3.1). RNA tag density tracks were generated from '.bam' files with RNA-STAR (2.4.2a) using '--outWigType bedGraph' and '--outWigStrand Unstranded' options for browser track viewing. Additional gene and transcript level quantification based on the GENCODE (v25) annotation was performed using featureCounts (http://bioinf.wehi.edu.au/featureCounts/) as contained in the Biocondutor Rsubread package (https://bioconductor.org/packages/release/bioc/html/Rsubread.html). Normalization and differential gene expression analysis between samples was conducted with EdgeR (v3.8). Specifically, 'featureCounts' were used for analysis and a threshold of median > 1 was applied to remove genes that were lowly/not expressed across samples. Subsequently, the following EdgeR functions were applied, in order, to samples within an experiment: DGEList, model.matrix, estimatedGLMCommonDisp, estimatedGLMTrendedDisp, estimatedGLMTagwiseDisp, and glmFit. Subsequently, the glmLRT function was applied to pairwise comparisons of singly or multiplex cells versus control cells to generate gene expression change P-values. Volcano plots were generated by plotting the resulting -log10(P-value) versus the log2(gene expression fold-change) for each group of singly or multiplex repressed cells versus control cells (no repressor RNA treated).

CUT&RUN data collection and analysis

CUT&RUN was performed as described[53]. Briefly, 4x10E5 CD3+ T cells were rinsed twice with wash buffer (20mM HEPES pH7.5, 150 mM NaCl, 0.5 mM spermidine) supplemented with proteinase inhibitor cocktail (Roche 4693159001) and incubated with 15 µl Concannavalin A-coated beads (Bangs Laboratories BP-531). The beads were set clear on a magnet and resuspended in 100 µl antibody binding buffer (20mM HEPES pH7.5, 150 mM NaCl, 0.5 mM spermidine, 0.02% digitonin, 2mM EDTA pH8) with a primary antibody and incubated overnight at 4C. The beads were set clear on a magnet and washed twice with digitonin-wash buffer (20mM HEPES pH7.5, 150 mM NaCl, 0.5 mM spermidine, 0.02% digitonin) followed by incubation with a secondary antibody (1:100 dilution in 100 µl digitonin-wash buffer) for an additional 1 hour at 4C and washed twice if the host of primary antibody is not rabbit. The beads were resuspended in 100 µl digitonin-wash buffer with 0.5 µl protein-MNase (final 14 µg/ml, kindly provide by Dr. Steven Henikoff from FredHutchinson Cancer Research Center) and incubated at 4C for an hour. The beads were set clear on a magnet and washed twice with digitoxin-wash buffer and then resuspended in 100 µl digitonin-wash buffer. 2 µl of 0.1 M CaCl2 were added to activate MNase and the digestion went on for 30 min at 0C. The digestion was stopped by adding 100 µl stop buffer (340 mM NaCl, 20 mM EDTA pH8, 4mM EGTA, 0.02% digitonin, 20 µg/ml glycogen, 50 µg/ml RNase A) and the reaction mixture was incubate at 37C for 10 min. The beads were spun at 16,000 g for 5 min at 4C and set clear on magnet. Supernatant was collected for proteinase K digestion (30C at 55C) and DNA were cleaned by phenol-chloroform-iso amyl alcohol extraction and ethanol precipitation. Antibodies used were obtained from following suppliers: FLAG (Sigma M2 1804), rabbit anti-mouse IgG (Jackson ImmunoResearch 31-005-048), H3K4me3 (Cell Signaling 9751), H3K9me3 (Abcam 8898). Sequencing libraries were generated with Takara's Thruplex DNA-Seq kit (R400677 +R400660), library QC\quantification was performed on our Fragment Analyzer (Agilent) using kit # DNF-474-1000 (High Sensitivity NGS). Sequencing was performed at 2x76bp on Illumina's Hiseq 4000. Raw sequencing reads were trimmed to remove adapter sequences and aligned to the human genome (hg38/GRCh38, no alts). Alignment was performed using bwa (version 0.7.12) with the following parameters: "-Y -l 32 -n 0.04" and "-n 10 -a 750" for alignment and mate-pairing (aln and sampe, respectively). Peak calls were generated with Hotspot (hotspot2; http://github.com/Altius/hotspot2). Read depth normalized signal, QC metrics were produced as described in https://www.encodeproject.org/pipelines/ENCPL202DNS/.

Immunofluorescence Imaging

T cells were debeaded, washed twice with PBS, and resuspended in 1x PBS for a final cell density of 1-2 million cells per ml. 30 ul of cell solution was then seeded into Poly-L-Lysine (PLL)-coated wells of a sterile 24-well plate (CellVis) and incubated for 20 minutes at RT. Cells were then fixed with 4% PFA (Polysciences Inc, #18814-10) in PBS for 10 minutes at room temperature. Fixed cells were washed 3x with 1x PBS, permeabilized with 0.5% Triton X-100 in PBS for 10 minutes at room temperature, blocked for 1 hour with 2% BSA (Jackson Immunoresearch, #001-000-161), and then incubated for 2 hours at room temperature with primary antibodies against FLAG

(F1804, mouse, 1:1000 dilution) in 2% BSA/1x PBS. Subsequently, cells were washed 3x with 0.05% Tween-20 (Bio-rad, #161-0781) in PBS, and then incubated for 1 hr with donkey anti-mouse Cy5 (1:500 dilution, #711-166-152, Jackson Labs) secondary antibody in 2% BSA/1x PBS. Lastly, cells were counterstained with DAPI (100ng/mL in 1x PBS) for 10 minutes at room temperature and washed 3 times with 0.05% Tween in PBS prior to mounting on glass slides using Prolong Gold (Molecular Probes P36930). Each wash step lasted 3 minutes at room temperature and was performed on a shaker (Stovall Inc). Samples were imaged using an inverted Nikon Eclipse Ti widefield microscope equipped with an Andor Zyla 4.2CL10 CMOS camera with a 4.2-megapixel sensor and 6.5μm pixel size (18.8mm diagonal FOV). Focused 2D cell images were acquired using a 60x Nikon Plan Apo 1.4 NA oil objective. Acquired images were subject to 100 rounds of iterative blind deconvolution using Microvolution software (Microvolution, CA) to minimize the effect of out-of-focus blurring that is inherent to widefield microscopy optics. Deconvolved images were processed using in-house Matlab (version 2017B, Mathworks, Natick, MA) scripts to numerically estimate the FLAG protein content in every cell nucleus, and for downstream statistical analysis

**References:**

51. T. Cermak *et al.*, Efficient design and assembly of custom TALEN and other TAL effector-based constructs for DNA targeting. *Nucleic Acids Res* **39**, e82 (2011).

52. T. Sakuma *et al.*, Efficient TALEN construction and evaluation methods for human cell and animal applications. *Genes Cells* **18**, 315-326 (2013).

53. P. J. Skene, J. G. Henikoff, S. Henikoff, Targeted in situ genome-wide profiling with high efficiency for low cell numbers. *Nat Protoc* **13**, 1006-1019 (2018).

**Fig. S1**

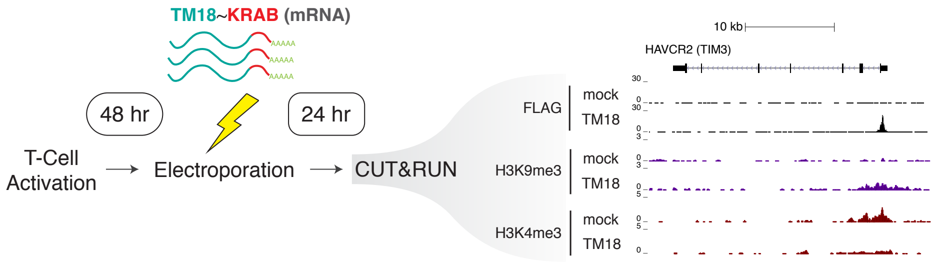


**Fig. S1. Synthetic repressors trigger chromatin state modification at target sites**. Activated T cells were electroporated with no RNA (mock) or the *TIM3* repressor TM18, and CUT&RUN was performed 24 hours later. Chromatin binding by TM18 at the *HAVCR2* (*TIM3*) locus was confirmed by CUT&RUN against the FLAG tag present on the TM18 repressor (right, black top panels showing sharp peak at the *TIM3* promoter). Downstream chromatin modifications were determined by CUT&RUN, confirming localized deposition of the repressive mark H3K9me3 (purple), and removal of the activity-associated mark H3K4me3 (red).

**Fig. S2**

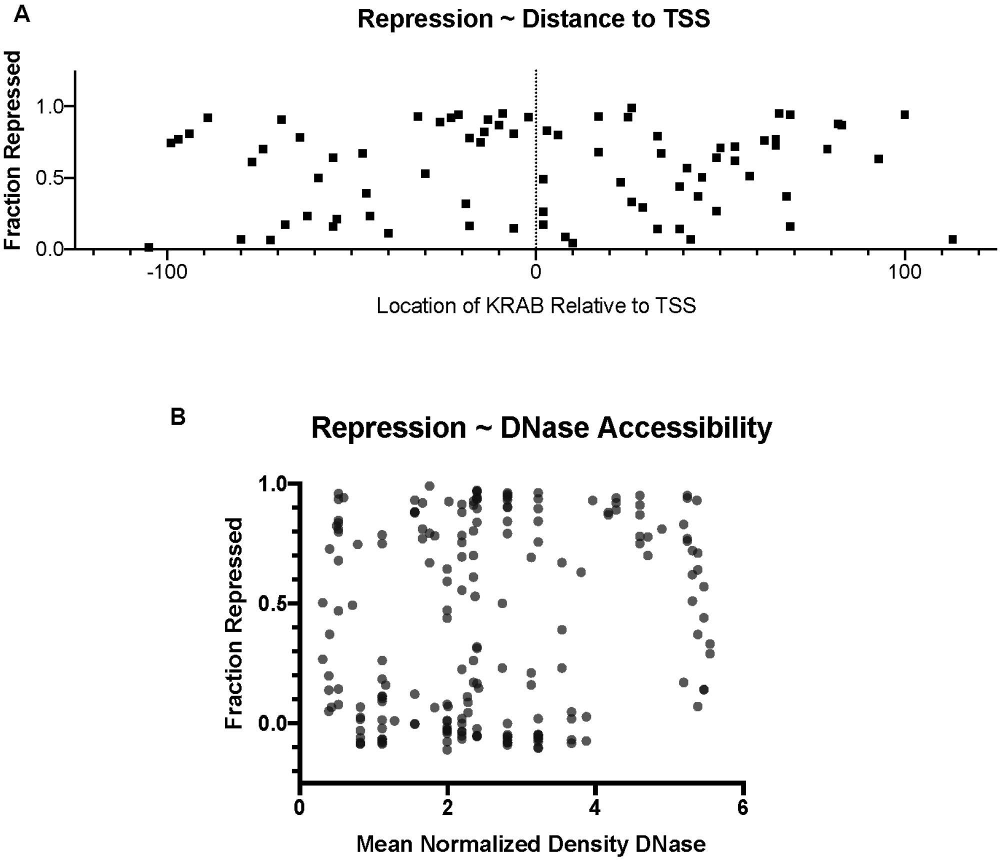


**Fig. S2. Synthetic repressor activity as a function of DNase accessibility at repressor binding site.** (a) Fraction of cells repressed (1 - fraction+ repressed / fraction+ control) is shown for all repressors targeting *PD-1*, *TIM3*, and *CTLA4* as a function of distance between the target gene's transcription start site (TSS) and the C-terminal RVD of the T-DBD, bearing the KRAB repressor domain. (b) Fraction repressed is shown for all repressors tested targeting *LAG3*, *TIM3*, *CTLA4*, and *PD-1* as a function the mean normalized DNase density of activated T-cells at the C-terminal RVD of the T-DBD, bearing the KRAB repressor domain, in a 10bp window.

**Fig. S3**

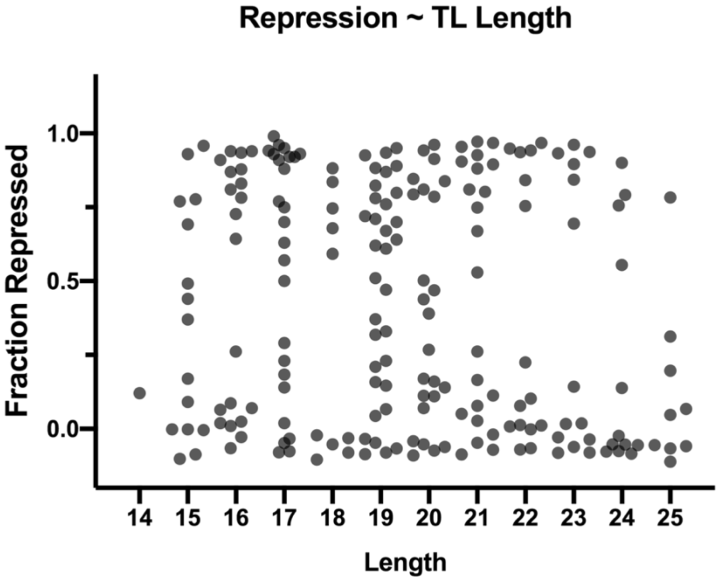


**Fig. S3. Synthetic repressor activity as a function of DBD length.** Fraction repressed (1 - fraction+ repressed / fraction+ control) is shown for all repressors tested targeting *LAG3*, *TIM3*, and *PD-1* as a function of number of repeat units in each DBD.

**Fig. S4**

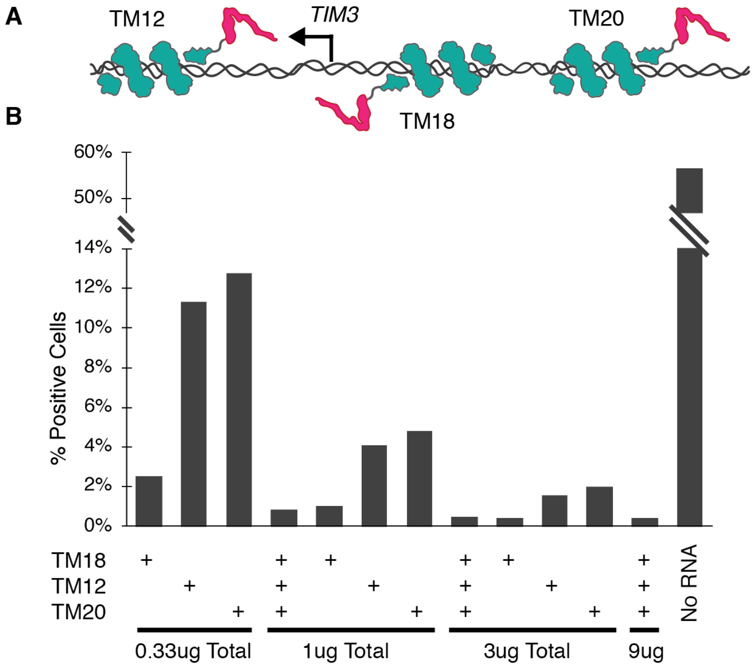


**Fig. S4. Synthetic repressor activity is dose-dependent and can be multiplexed at an individual promoter to quantitatively modulate repression**. (a) Experimental diagram showing the relative positions of three T-DBD-KRABs targeting *TIM3* used to test dose dependence and effect of multiplexing at a single promoter. (b) Percent *TIM3*-positive cells at 48 hours post-transfection with the indicated amounts of the indicated T-DBD-KRAB combinations was measured by flow cytometry. Multiplexing repressors at a single promoter can achieve better than additive effects.

Fig. S5

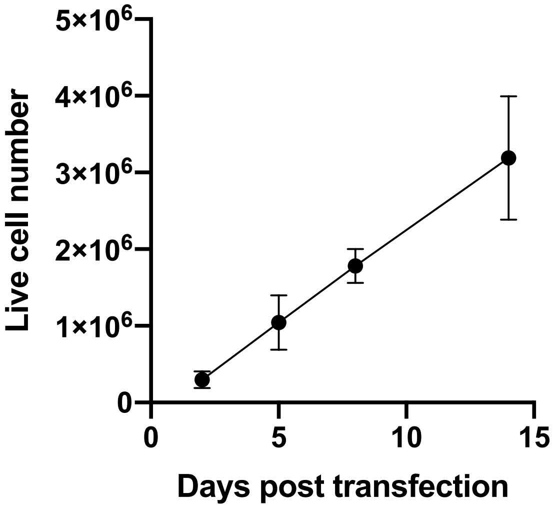


**Fig. S5. Expansion of T cells after activation and transfection.** Error bars represent standard deviation of four individual wells in a single experiment. Population doubling time was calculated assuming a constant proliferation rate and cell counts at days 2 and 5.

**Table S1:** Synthetic repressor target sequences

| Name | Target sequence (5'-3') | RVD sequence (N-C) |
|---|---|---|
| TM18 | TGGCAGTGTTACTATAA | NH-NH-HD-NI-NH-NG-NH-NG-NG-NI-HD-NG-NI-NG-NI-NI |
| PD02 | TGGTGGGGCTGCTCC | NH-NH-NG-NH-NH-NH-NH-HD-NG-NH-HD-NG-HD-HD |
| LG09 | TGCCGTTCTGCTGGTCT | NH-HD-HD-NH-NG-NG-HD-NG-NH-HD-NG-NH-NH-NG-HD-NG |
| TM12 | TGGCAATCAGACACCCGGGTG | NH-NH-HD-NI-NI-NG-HD-NI-NH-HI-HD-NI-HD-HD-HD-NH-NH-NH-NG-NH |
| TM20 | TGCCACACTACACACAT | NH-HD-HD-NI-HD-NI-HD-NG-NI-HD-NI-HD-NI-HD-NI-NG |
